# Dynamics of central remyelination and treatment evolution in a model of Multiple Sclerosis with Optic Coherence Tomography

**DOI:** 10.1101/2020.10.22.350181

**Authors:** Rocío Benítez-Fernández, Carolina Melero-Jerez, Carmen Gil, Enrique de la Rosa, Ana Martínez, Fernando de Castro

**Author notes:** These authors contribute equally to the work (Rocío Benítez-Fernández, Carolina Melero-Jerez). Correspondence: Prof. Ana Martinez, Centro de Investigaciones Biologicas-CSIC, Ramiro de Maeztu 9, 28040 Madrid (Spain), Dr. Fernando de Castro, Instituto Cajal-CSIC, Avda. Doctor Arce 37, 28002 Madrid.

## Abstract

The need for remyelinating drugs is essential for healing important diseases such as multiple sclerosis (MS). One of the reasons for the lack of this class of therapies is the impossibility to follow remyelination *in vivo*, which is of utmost importance to perform good clinical trials. Here, we show how the optical coherence tomography (OCT), a cheap and non-invasive technique commonly used in ophthalmology, may be used to follow remyelination *in vivo* in MS patients. Our pioneer study validates the study of myelin/remyelination in the optic nerve using OCT and reflects what is occurring in non accessible CNS structures, like the spinal cord. For this study we used the oral bioavailable small molecule VP3.15, confirming its therapeutical potential as neuroprotective, antinflammatory and remyelinating drug for MS. Altogether, our present results confirm the usefulness of OCT to monitor the effectivity of remyelinating therapies *in vivo* and underscore the relevance of VP3.15 as potential disease modifying drug for MS therapy.

## Introduction

Multiple sclerosis (MS) is the most common primary demyelinating disease and neurological condition affecting young adults, with about 2.5 million people currently diagnosed with MS around the world (1). It is characterized by glial cell pathology (especially oligodendrocytes and their precursors), demyelination, inflammatory processes and axonal damage in the central nervous system (CNS) (2). In addition to glial cell pathology, the destruction of CNS myelin is accompanied by the activation of macrophages and microglia, cells present in MS lesions and behave either as proinflammatory or anti-inflammatory agents depending on the stage of the lesion (3–6).

An unmet challenge in demyelinating diseases like MS is reestablishing the lost myelin, thus reducing the neurological dysfunction. A proper approach should include three steps: i) the modulation of the inflammatory response; ii) the protection of oligodendrocytes; and iii) the promotion of effective remyelination. Current available treatments are exclusively immunomodulatory, focused on the reduction of inflammation, which ameliorates the evolution of the non-progressive forms of the disease but does not cure them (7). During the last decade, many studies have investigated new myelin repairing/regenerating mechanisms. To date, only clemastine fumarate, an old-fashioned antihistaminic drug, has shown relative efficiency on randomized controlled clinical trials as a remyelinating drug (8, 9). Many other targets have been explored with this aim, such as the pathways of Notch and Wnt signaling (10), glutamatergic receptors (11), nuclear receptors such as RXRy, PPARy and VDR (12–14), and components of the extracellular matrix (15, 16). Furthermore, different growth factors and chemotropic molecules, like FGF-2 and Anosmin-1, secreted semaphorins and others, have been suggested to cause spontaneous remyelination in human MS (17, 18). However, the efficacy of remyelinating therapies is difficult to assess *in vivo* due to the technical limitations of accessing the CNS from the outside, not only in clinical trials but also in preclinical studies where MS animal models were used.

Consequently, there is a need for novel tools to real-time measure and quantify the myelin loss and the effective remyelination after a potential regenerative treatment. Over the past few decades, advanced visualization techniques such as 7-Tesla Magnetic Resonance Imaging (MRI), Positron Emission Tomography (PET) or visual evoked potentials have become pivotal in the diagnosis and monitoring of MS. They have allowed the study of its different aspects, such as the perivascular inflammation, the bursting of new CNS lesions, the activation of the immune response and the inflammation of leptomeninges (19). However, because clinical MS reflects a combination of inflammation, degeneration and regeneration, these techniques fail to detect the remyelination process (20). For this reason, the spectral domain Optic Coherence Tomography (OCT), a new imaging technology, has emerged in the last few years. OCT is a reproducible and non-invasive method for retinal structure visualization in physiopathological conditions, including neuroinflammatory disorders such as neuromyelitis optica (NMO) and optic neuritis (ON), both in animal models (21) and humans (22, 23). In patients, OCT measurements include the macula volume and the area and thickness of the retinal nerve fiber layer (RNFL), which comprises the axons of the retinal ganglion neurons that converge to form the optic nerve (24). Because retinal alterations have been described as a component of several MS outcomes, OCT use has been extended to MS in clinical practice for diagnostic purposes, although if the optic nerve reflects the situation in the parenchimatous CNS remains to be address (25–31). Although a correlation has been described between the decrease in RNFL thickness and brain atrophy in MS (32, 33), it has yet to be ascertained whether the results in the RNFL clearly compile the histopathology of the CNS, in terms of inflammation, demyelination status and oligodendrocyte-lineage cell availability and maturity. The finding of an accurate correlation between the myelin alterations in deep CNS structures and peripheral and technically approachable areas such as the retina would improve the research into remyelinating drugs, elucidating their real-time efficiency in a quantitative manner. It should be noted that, in patients, it is easier to visualize the structure of the retina and the optic nerve head taking advantage of imaging techniques, but it is more difficult to establish correlations within deeper parts of the CNS; hence the need to validate the OCT in animal models.

The main goal of the present study isto validate OCT as a methodology for the accurate follow-up of the neurological symptoms and the histopathological hallmarks in a murine model of MS, i.e experimental autoimmune encephalomyelitis (EAE). The second aim is to evaluate the remyelination efficacy of a dual phosphodiesterase-7 (PDE7) and glycogen synthase kinase-3β (GSK3β) inhibitor named VP3.15 (34). This compound is a small heterocyclic drug-like molecule with a high safety profile and good pharmacodynamic and pharmacokinetic properties following intraperitoneal administration (34). Furthermore, VP3.15 has been described as an inducer of remyelination *in vitro* and *in vivo* in oligodendrocyte precursor cells (OPCs) isolated from non-tumoural biopsies of human adult brain cortex (35, 36). This multitarget compound has also shown anti-inflammatory properties *in vivo*, both in retinal dystrophies (37) and in the EAE model of MS (38). The data presented supports OCT as a promising and and efficacious tool to evaluate the effects of remyelinating drugs in unobservable areas of the CNS. Moreover, a clear correlation between OCT data (retina, the papilla of the optic nerve) and immunohistochemical observations (optic nerve and spinal cord) is demonstrated. Finally, our results confirm the compound VP 3.15 as a promising remyelinating drug that merits further development into clinical trials.

## Results

### The PDE7/GSK3 dual inhibitor VP3.15 ameliorates clinical course of EAE

The dual inhibitor VP3.15, as effective as the oral FDA-approved drug fingolimod in EAE model (38) and with remyelating effect *in vitro, ex vivo and in vivo* (39), was the pharmacological tool chosen to test our hypothesis on the usefulness of OCT to monitor remyelination. Benefits of the repeated treatment with VP3.15 (see Methods) in EAE mice were evident since the beginning, with the clinical score (CS) decaying faster and to a significantly lower value in the VP3.15 treated EAE mice by day 19 after onset (Fig. 1b). For a more quantitatve estimation of the healing effect of VP3.15, we fitted the results to an exponential expression (40), and the score fraction (Sf) obtained was significantly better (0.2) than the EAE-Vehicle group (0.3), thus the treatment with VP3.15 ameliorated EAE effects by 33% compared to the vehicle-treated mice (Fig. 1c) by the end of the experiment. In addition to that, the estimated recovery rate (ν) was higher in the treated animals (EAE-Veh= 0.187 days^-1^; EAE-VP3.15=0.226 days^-1^), what was reflected in the fact that EAE-VP3.15 reached the endpoint Sf of EAE-VEH nearly 10 days before.

**Fig 1.**
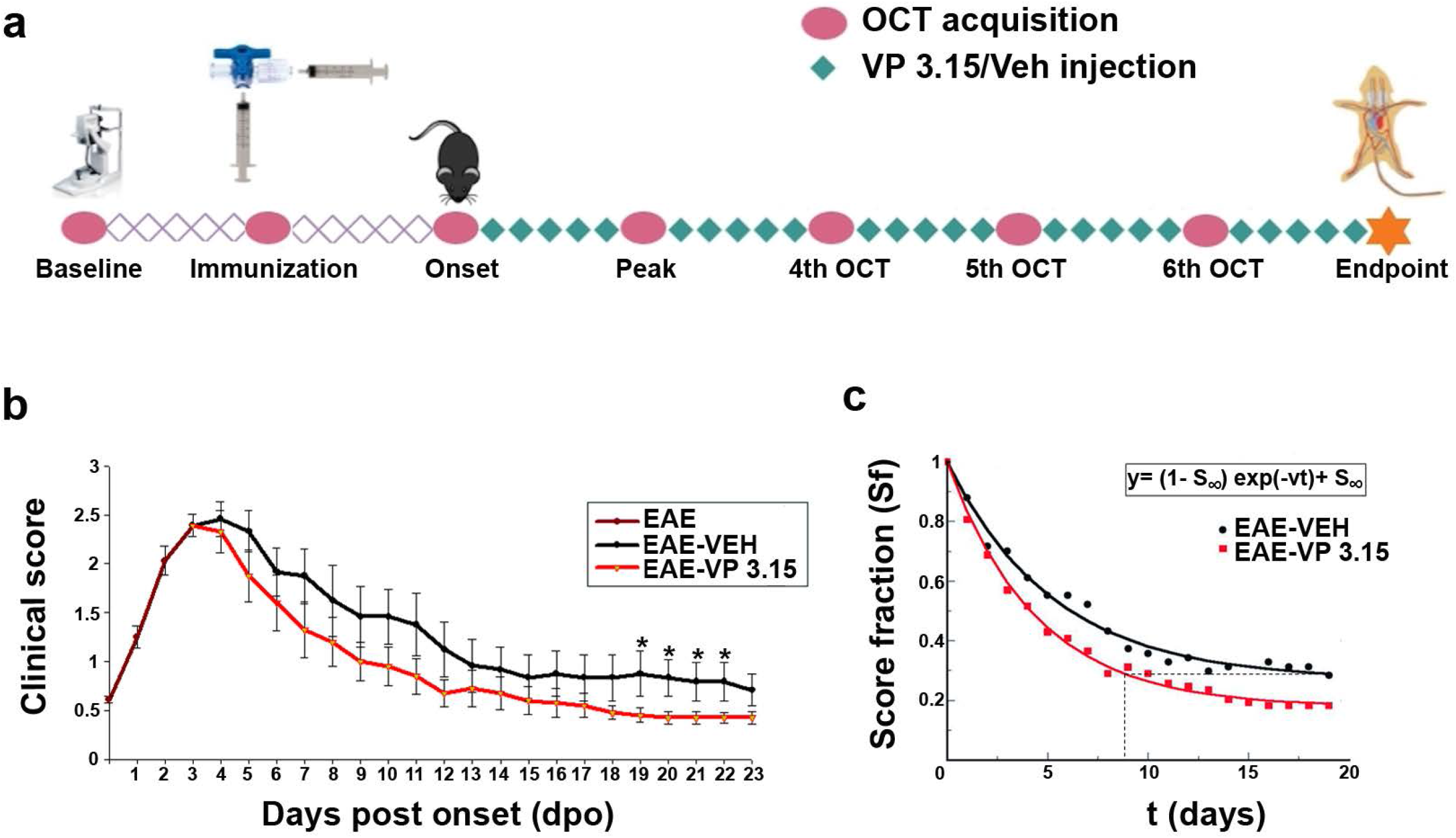
The treatment of EAE mice with the dual inhibitor VP3.15 ameliorates the clinical course from the beginning of the administration; **a**: Procedure outline; **b**: Time course representation of the clinical performance of EAE mice from the onset of symptoms, showing a lower clinical score in the EAE-VP3.15 compared to EAE-VEH (two-way ANOVA to compare the two treatments: p< 0.05; results of Student’s *t*-test are represented as: *p<0.05 from day 19 to day 22); **c**: Exponential expression of the score decay from the beginning of the VP 3.15 treatment, where the score (S) at every time point is normalized by the maximal score (Smax). The expression shows that VP3.15 maintains the score in a lower level than vehicle at longer times. Abbreviations: dpo= days post onset, Sf= score fraction; S∞= score at infinite times

### Retinal and optic nerve changes can be monitored using Optic Coherence Tomography in the EAE model

Nowadays, retinal layer shrinkage observed by OCT is used in clinics to predict terms of cognitive decline and brain atrophy in MS patients (33, 41, 42). Our main aim in this work was to check if remyelination, specifically, can be effectively monitored in real-time using a non-invasive technique such as the OCT (see Methods). Besides the predicable stability of retinal thickness in the SHAM group along the entire experiment, we observed that the retina was significantly thinner when EAE was induced. As expected the treatment with VP3.15 showed better dynamics than EAE-VEH, including significant recovery on the sixth OCT onwards (Fig. 2 e). These results were more consistent when we measured the optic nerve width: from fourth OCT to endpoint, the group treated with VP3.15 showed significantly largest OCT measures than non treated EAE one (Fig. 2f; results of Two way-ANOVA Bonferroni *post-hoc* test were p<0.05). Concerning the retina, there was a significant recovery of VP3.15 treated animals after the fifth OCT and maintained at that recovered the SHAM-like levels at the endpoint. To further assess the remyelination of the damaged tissue in the EAE mice, we performed a detailed tissue analysis of the optic nerve (rostral localization, thus closer to the eye) and spinal cord (caudal localization) obtained at the endpoint.

**Fig 2.**
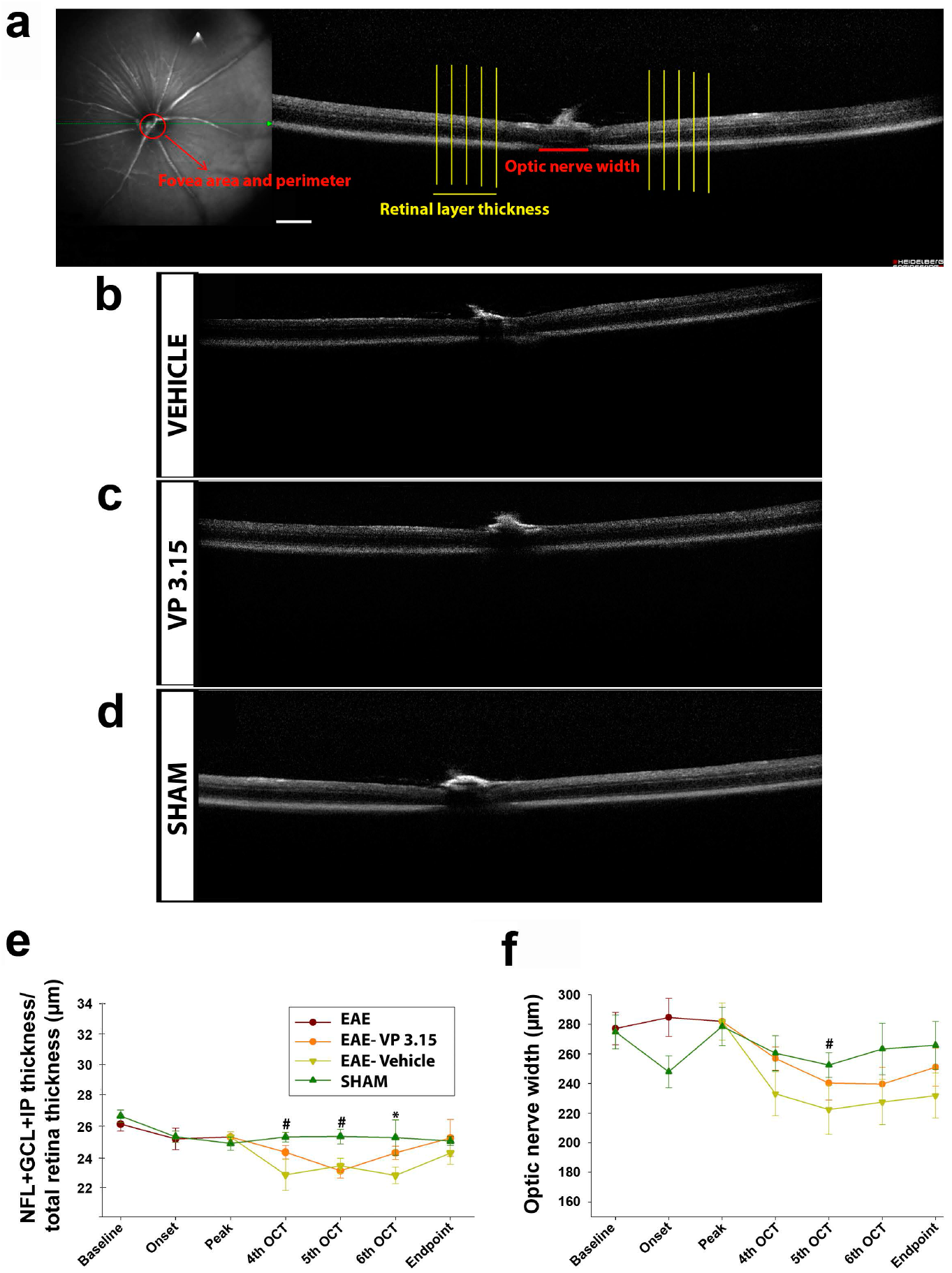
Retina and optic nerve analysis by OCT of the EAE mice reveals changes associated to the concurrent disease; **a**: Schematic representation of the different measurements at the fovea, optic disc and retina levels; **b-d**: Representative images obtained with the OCT; **e**: Retinal layer thickness analysis along the experiment; **f**: Dynamics of optic nerve width along the experiment. Scale bar represents 200 µm in a-d. Abbreviations: GCL= ganglion cell layer; IPL= inner plexiform layer; NFL= nerve fiber layer. Results of Student’s *t*-test are represented as: *p<0.05 (EAE-VP3.15 vs. EAE-VEH) or #p<0.05 (EAE-VEH vs. SHAM)

### The remyelinating role and the effect in the oligodendrocytes lineage of VP 3.15

We checked at endpoint the histopathology of two out of the most myelinated structures in the CNS, the optic nerve and the spinal cord. In the optic nerve, the treatment with VP3.15 showed an increase higher than 50% in myelinated (MBP^+^) axons when compared with EAE-VEH (Fig. 3A, c-d, and f-g). In addition, the integrity of the optic nerve axons (NFH^+^) was better preserved in the VP3.15-treated animals compared to the EAE-VEH group (Fig. 3 b, e and h). Similar results were obtained in the spinal cord: the demyelinated area in the EAE-VP3.15 group was significantly lower and showed more MBP^+^ staining than in the EAE-VEH (Fig. 4a-d, g-h and i-j), while the axons were also better preserved in the VP3.15-treated group (Fig. 4e-f and k). Although it is difficult to compare both structures, our results in the optic nerve seemed slightly more robust than in the spinal cord (percentages in Fig. 3g-h *vs* in Fig. 4i-k), and the remarkable differences in myelination and axonal density could explained them. These structural differences were observed in EAE-VEH and also when the treatment with VP3.15 was applied, growing both parameters (Fig. 5a-b). Altogether, present results confirm previous data obtained in different demyelinating animal models of MS, including EAE, and *in vitro* (35, 36). In addition to that, we found a direct and significant correlation between the normalized MBP area and the normalized axonal area in both treatments, both in the optic nerve (Fig. 5c; EAE-Veh: r=0.559, p>0.01; EAE-VP3.15: r=0.546, p>0.01) and the spinal cord (Fig. 5d; EAE-Veh: r=0.933, p>0.001; EAE-VP3.15: r=0.803, p>0.001).

**Fig 3.**
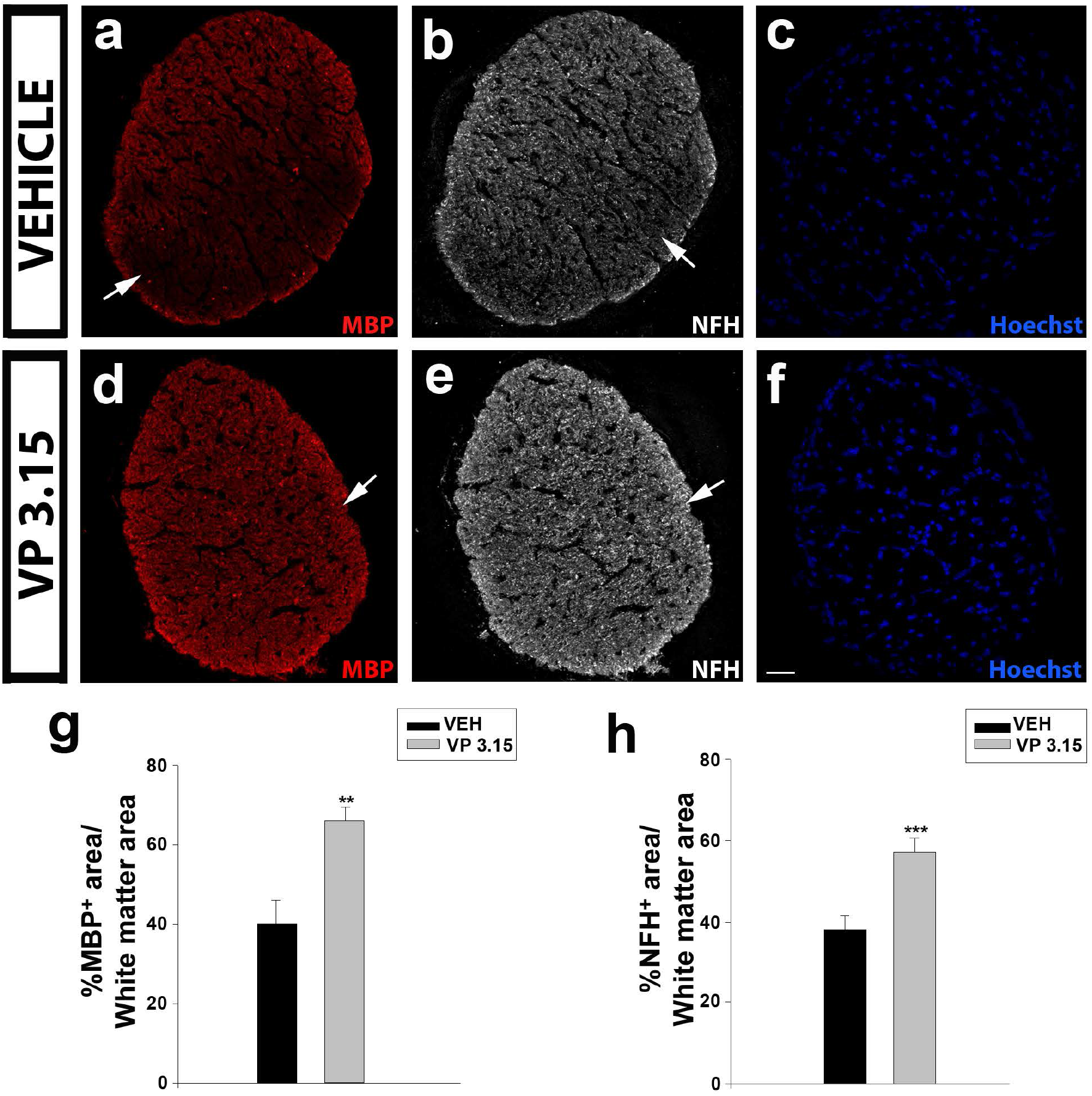
The VP 3.15-treated mice preserve the optic nerve from myelin loss and axonal damage; **a-f**: Detailed views of the optic nerve of vehicle (a-c) and VP3.15-treated mice (d-f), labelled for MBP (red), NFH (grey) and nuclei (Hoechst staining; blue); **g-h**: Histograms showing a significant increase in the percentage of both MBP^+^ (j) and NFH^+^ (k) area in the VP3.15-treated compared to the vehicle-treated mice. Scale bar represents 50 µm in a-f. Abbreviations: MBP= myelin basic protein; NFH= neurofilament heavy. Results of Student’s *t*-test are represented as: ** p< 0.01; ***p < 0.001

**Fig 4.**
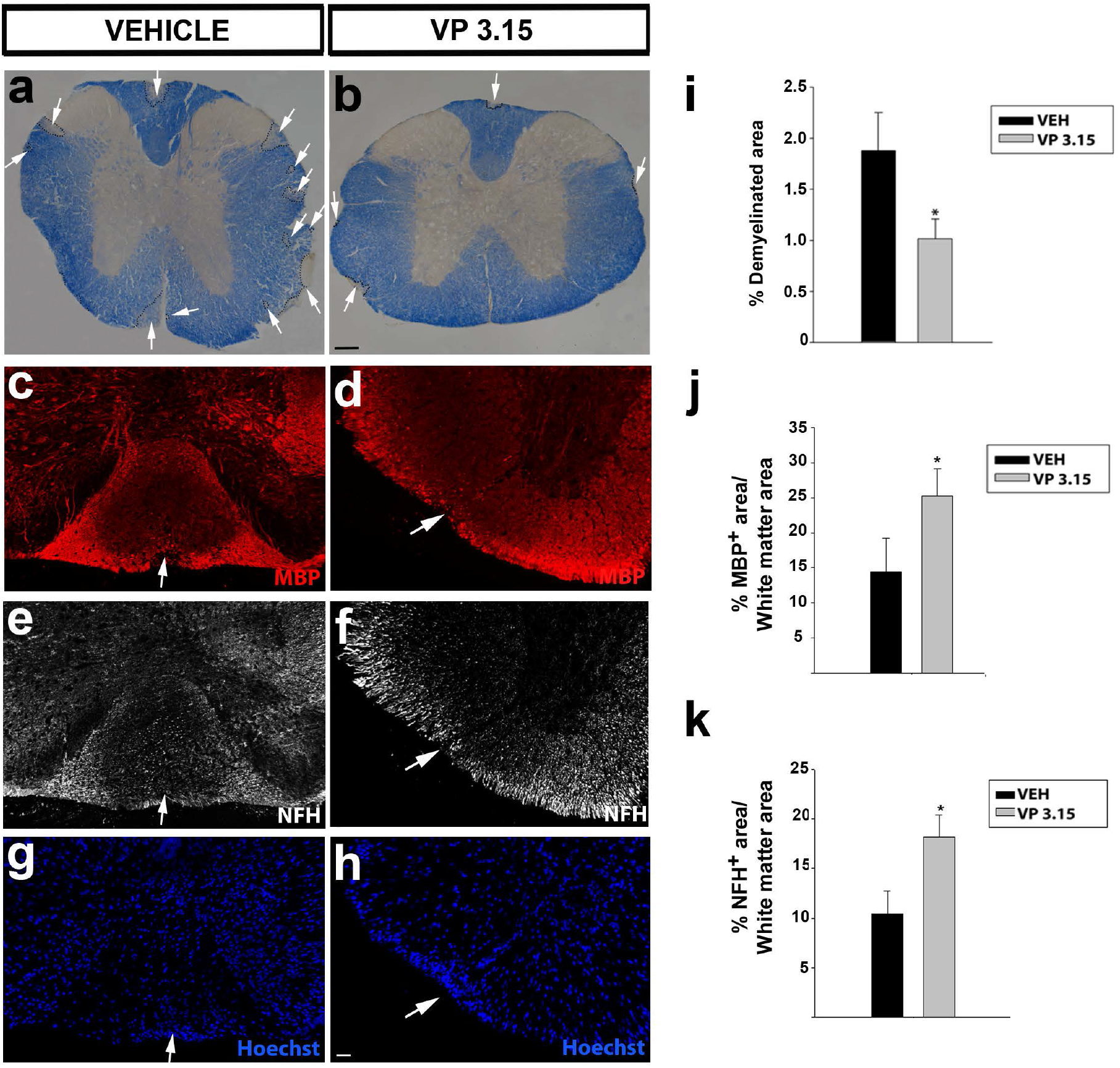
The spinal cord of VP3.15-treated mice presented a lower level of demyelination and axonal damage; **a-b**: Panoramic views of the spinal cord stained with eriochrome cyanine of vehicle (a) and VP3.15-treated mice (b). The demyelinated area is delimited by the dashed line and arrows. Scale bar represents 200 µm in a-b; **c-h**: Detailed views of the spinal cords of vehicle (left) and VP3.15-treated mice (right), labelled for MBP (red), NFH (grey) and nuclei (Hoechst staining; blue). Lesions are identified as an accumulation of nuclei; **i**: Graph showing a decreased percentage of demyelinated area respect to the white area in the VP3.15-treated mice; **j-k**: Histograms showing a significant increase in the percentage of both MBP^+^ (j) and NFH^+^ (k) area in the VP3.15-treated compared to the vehicle-treated mice. Scale bar represents 50 µm in c-h. Abbreviations: MBP= myelin basic protein; NFH= neurofilament heavy. Results of Student’s *t*-test are represented as: * p< 0.05

**Fig 5.**
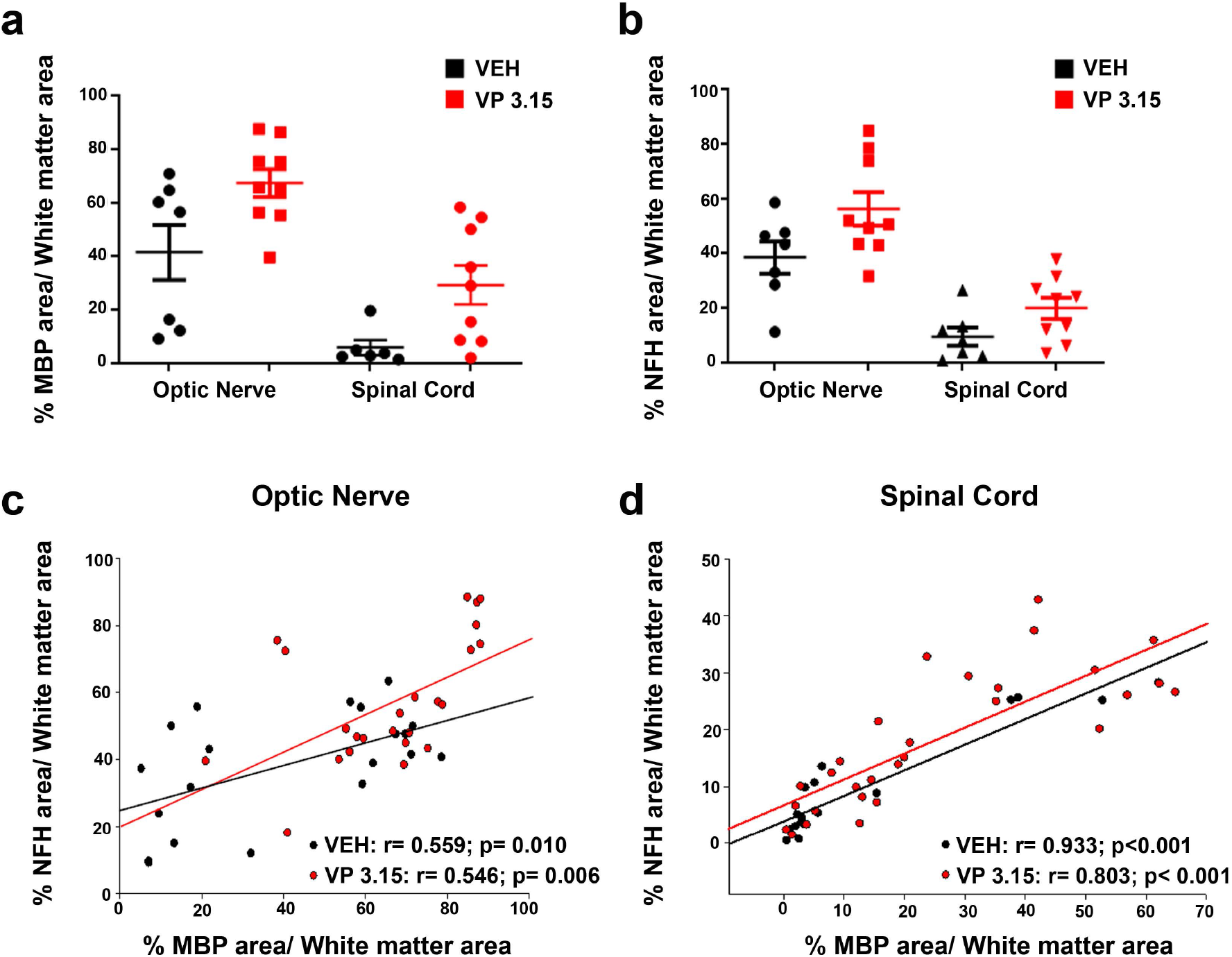
Axonal integrity is higher when the myelin is better preserved**; a-b**: Graphs showing the percentage of MBP (a) and NFH (b) area respect to the total white matter area between structures; **c-d:** Graphs showing the results of Pearson’s correlation tests between the normalized MBP area and the normalized axonal area among treatments and in the optic nerve (c) and spinal cord (d)

Also, it is known that the dual PDE7-GSK3 inhibition by VP3.15 enhances murine and adult human OPC differentiation without affecting their survival or proliferation (35). We inquired whether the remyelinating role and the neuroprotection on axons already seen could be related to changes in the oligodendrocytes lineage. In the optic nerve, a significant increase in the number of precursor cells, labelled as PDGFRα^+^ cells (Fig 6a, e and i) and mature cells, identified as CC1^+^ cells (Fig 6c, g and j) was observed in the VP3.15 treated animals compared to the vehicle group. In the spinal cord, the same effect, an increase in the precursor cells (Fig 7 a, c, g, i and m) and mature oligodendrocytes (Fig 7b, e, h, k and n) was also observed. At this respect, we applied a two way ANOVA to unreavel the differences among treatments and structures, showing no significant differences for PDGFRα^+^ cells (Fig. 8A; p= 0.072). We did found a significant difference between structures in the case of CC1^+^ cells, being the proportion lower in the optic nerve than in the spinal cord (Fig. 8d; p<0.001). We then looked for a relationship between the myelin (MBP^+^ cells) and the two stages of the oligodendrocyte lineage under study (Fig. 8b, c, e and f). We found a significant and direct correlation in the case of CC1^+^ cells in the optic nerve of vehicle-treated EAE mice, (Fig. 8b and E; EAE-Veh: r= 0.470, p= 0.0422; EAE-VP3.15: r=0.254, p= 0,22). The relation between a lower proportion of CC1^+^ cells and higher levels myelination that we previously showed could be explained by a difference in the rate of differentiation to mature phenotypes, that might be higher in the case of a highly myelinated tissue as is the optic nerve.

**Fig 6.**
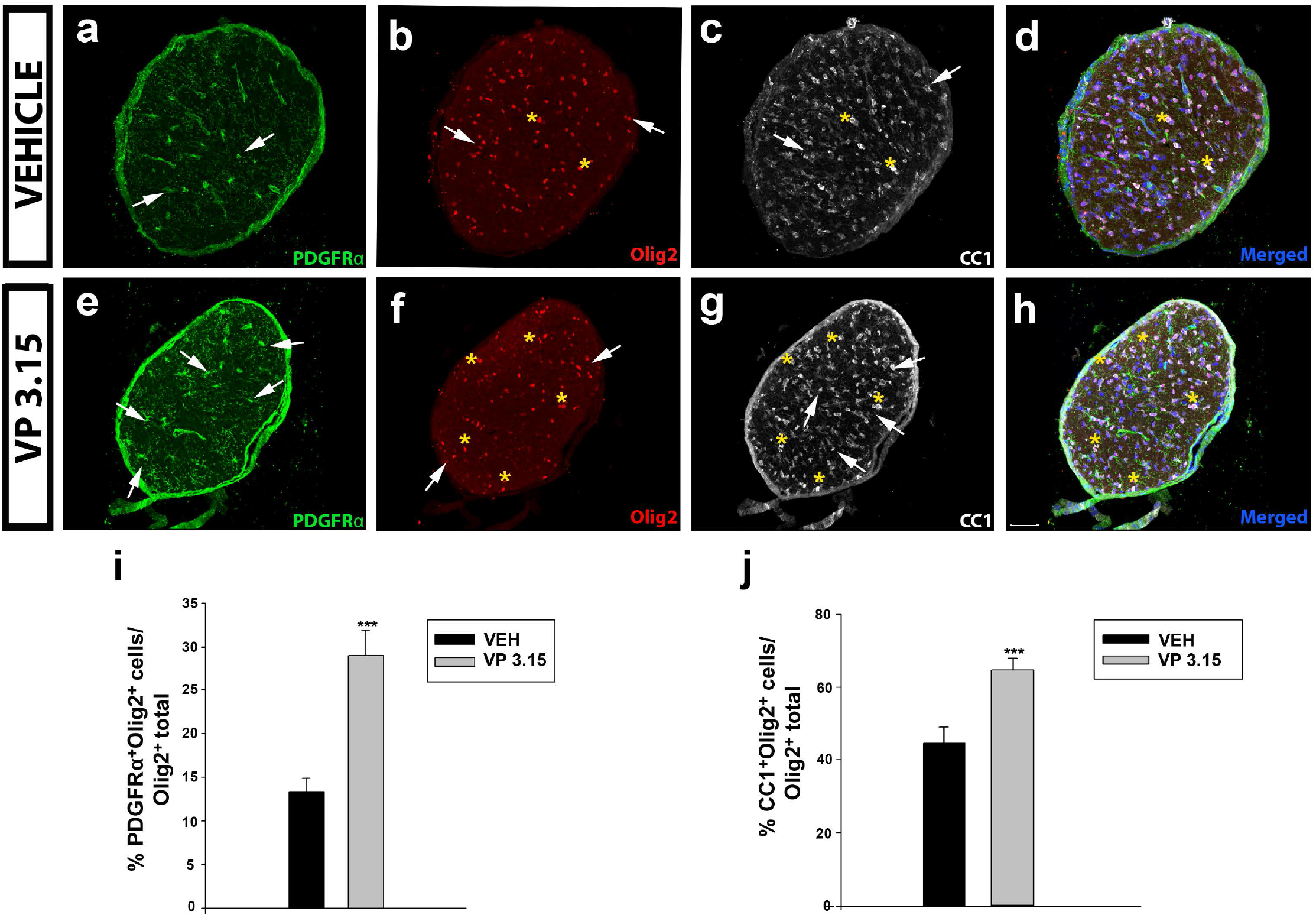
The VP 3.15 promotes the presence of both precursor and mature oligodendrocytes in optic nerve; **a-h**: Detailed views of the optic nerve of vehicle (a-d) and VP3.15-treated mice (e-h), labelled for PDGFRα (green), Olig2 (red), CC1 (grey) and merged includes nuclei (Hoechst; blue). Arrows point to PDGFRα^+^Olig2^+^ cells and asterisks to CC1^+^Olig2^+^ cells. Scale bar represents 50 µm in a-h; **i-j:** Graphs showing the significant increase in the percentage of PDGFRα^+^Olig2^+^ cells (i) and CC1^+^ Olig2^+^ cells (j) after the treatment with VP 3.15 compared to the vehicle. Results of Student’s t-test are represented as: ***p < 0.001

**Fig 7.**
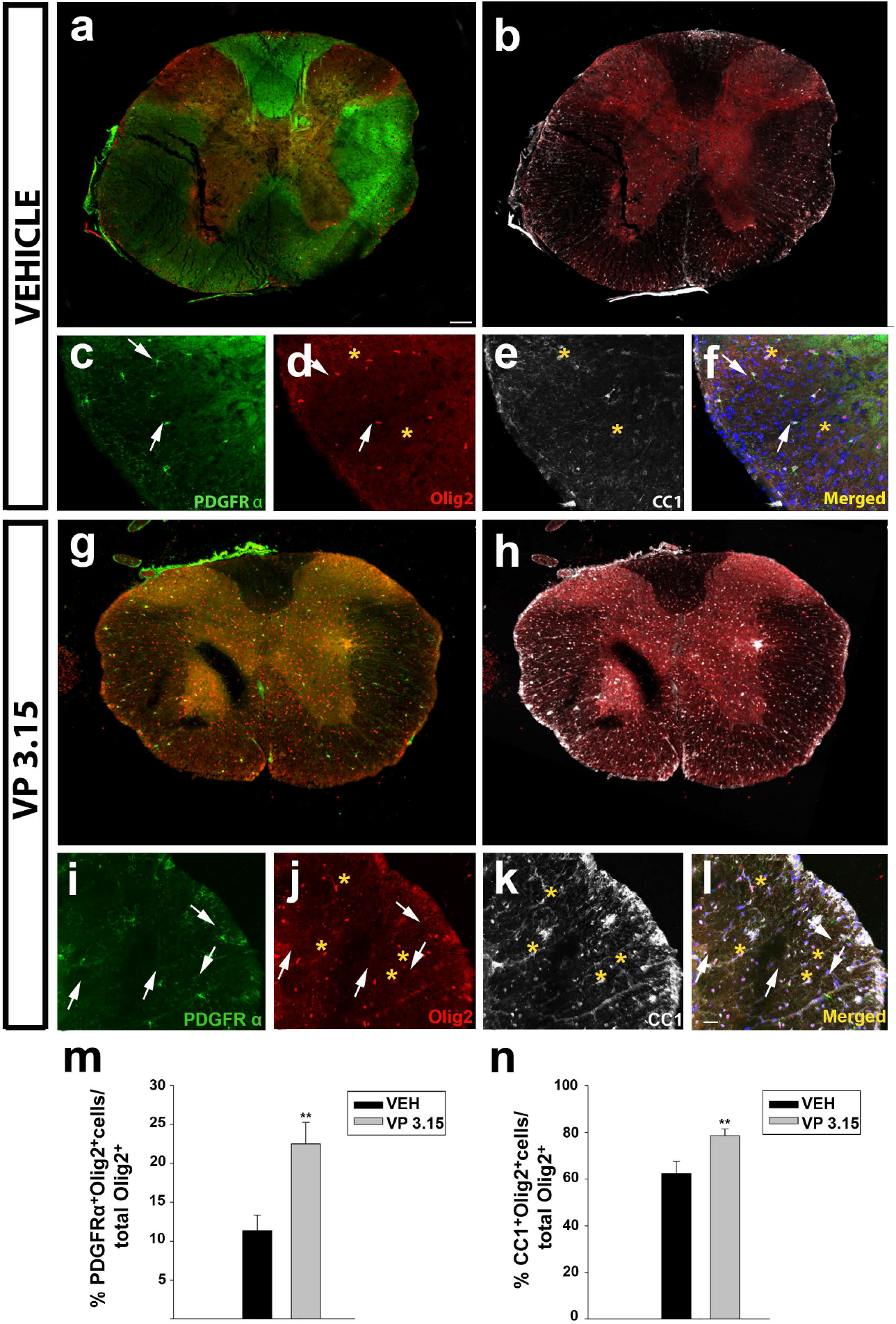
The treatment of EAE mice with VP 3.15 promotes the presence of both precursor and mature oligodendrocytes in spinal cord; **a-l**: Representative panoramic views (a-b and g-h) and detailed images (c-f and i-l) of the spinal cords of vehicle (a-f) or VP 3.15-treatment (g-l). Cells are labelled for PDGFRα (green), Olig2 (red), CC1 (grey) and merged, that include nuclei (Hoechst; blue). Arrows point to PDGFRα^+^Olig2^+^ cells, and asterisks to CC1^+^ Olig2^+^ cells. Scale bar in a, b, g and h represents 200 µm and the rest represents 50 µm; **m-n**: Graphs showing the significant increase in the percentage of PDGFRα^+^Olig2^+^ cells (m) and Olig2^+^ CC1^+^ cells (n) after the treatment with VP 3.15 compared to the vehicle. The sample size was: EAE-VEH= 7 mice; EAE-VP3.15= 10 mice. Results of Student’s *t*-test are represented as: **p < 0.01

**Fig 8.**
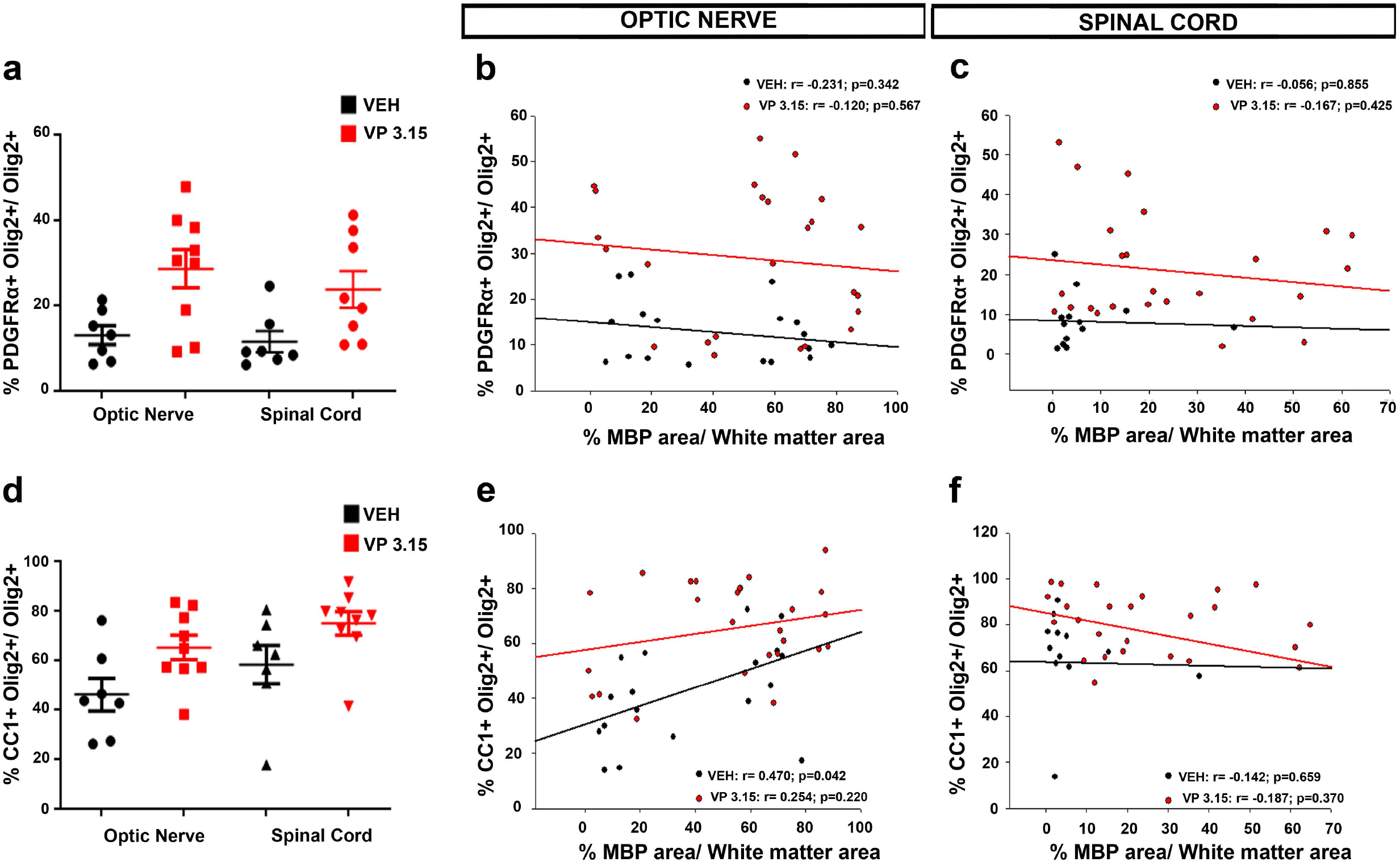
The maturation state of oligodendrocytes is related to the myelin preservance in the optic nerve; **a-d:** Graphs showing the different levels of precursor (PDGFRα^+^; a) and mature cells (CC1^+^ cells; d) between structures; **b-c**: Graphs showing the correlation between the normalized MBP area and the percentage of PDGFRα^+^Olig2^+^ cells in both treatments and in the optic nerve (b) and spinal cord (c); **e-f:** Graphs showing the results of Pearson’s correlation between the normalized MBP area and the CC1+ Olig2^+^ cells in both treatments and in the optic nerve (e) and spinal cord (f)

### VP3.15 treatment modifies microglial activation state

To get a deeper insight into the inflammatory component of the CNS at the end of the VP3.15 treatment, we analyzed the composition of the microglial population, being divided into three groups: ramified, stellate-shaped and amoeboid microglia (43, 44). In the optic nerve, there was an increase in the proportion of the stellate-shaped cells in the EAE-VP3.15 group compared to vehicle (Fig. 9 a, b and e; EAE-VEH= 0.043 ± 0.007 %; EAE-VP 3.15 = 0.082 ± 0.006 %), while the other two subpopulations remained similar between groups (Fig. 9 a, b and e; ramified microglia: EAE-VEH= 0.004 ± 0.010%; EAE-VP3.15= 0.045 ± 0.006%; amoeboid microglia: EAE-VEH= 0.044 ± 0.0147%; EAE-VP3.15= 0.027% ± 0.014%). Regarding the spinal cord, the microglial population was enriched in stellate-shaped cells in the VP3.15-treated mice in contrast to the vehicle-treated group (Fig. 9 c-d and f; EAE-VEH= 0.025 ± 0.003%; EAE-VP3.15= 0.045 ± 0.002%), being the amoeboid subpopulation significantly decreased (Fig. 9 c-d and f; EAE-VEH= 0.004 ± 0.003%; EAE-VP3.15= 0.266 ± 0.003%). Finally, that of the ramified microglia remained constant in both groups, as in the optic nerve. Altogether, our present results show that the remyelinating role of VP3.15 is accompanied by a switch of the microglial content to anti-inflammatory phenotypes.

**Fig 9.**
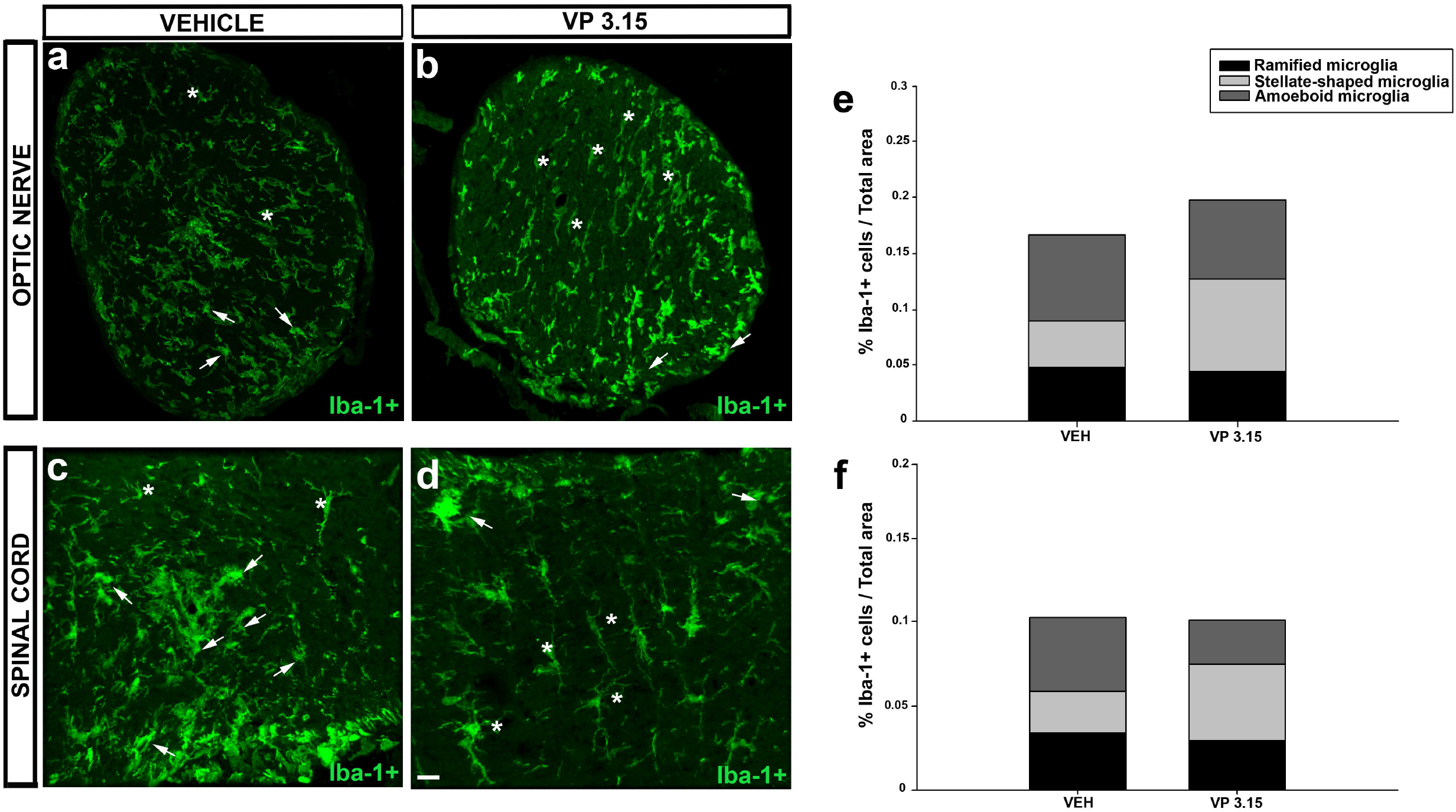
Effects of VP 3.15 on Iba-1^+^ in spinal cord and optic nerve; **a-d**: Detailed views of the optic nerve of vehicle (a) and VP 3.15-treated mice (b) and the spinal cord of vehicle (c) and VP3.15-treated mice (d), labelled for Iba-1^+^ cells (green). Arrows point to amoeboid microglia cells and asterisks mark stellated-shaped microglial cells; **e-f**: Stacked histograms of the spinal cord (e) and optic nerve (f) content in microglia. In both cases, whereas the ramified microglia remain constant in the EAE-VP3.15 and the EAE-VEH group, the proportion of a stellated-shaped and amoeboid microglia is the opposite, being higher in the former. Scale bar represents 50 µm in a-d

## Discussion

In the general frame of the unmet challenge for neurology to develop effective remyelinating approaches, our present study adds new evidences on the usefulness of OCT for monitoring MS evolution at least with remyelinating treatments. OCT is a non invasive technique with an easy translation to the patients (45) and data here presented show that it might be considered as an useful technique to follow remyelinization in patients under some treatment, mainly if it modulates myelin quantity. Furthermore, we have provided more details on the potential of VP3.15 as a useful drug candidate for the future therapy of MS.

To the best of our knowledge, this is the first study correlating the morphological characteristics of optic nerve *in vivo* with demyelination/neurodegeneration and remyelination in an animal model of MS. We corroborated the data that we obtained with OCT with the histological study performed at the endpoint of the study. Part of our findings, like the decreased thickness of RNFL and GCL have been described in MS (46–48), but neither that study was repeated at different times of evolution, nor it was corroborated with *postmortem* observations, two aspects that confer to our present study part of its uniqueness in the moment of its publication. We additionally propose that, in spite of the limitations of its observation via OCT, the width of the optic nerve papilla would be very effective to monitor the demyelination/remyelination dynamics of the entire optic nerve and the parenchymatous CNS, as seen by the multiple correlation found in our study. Given the parallel between the degree of myelin preservation/regeneration and the protection of axons observed in our current study, we propose that OCT could be also useful to monitor the overall neurodegeneration of neurons in the CNS. Techniques allowing the visualization of glia in response to drug discovery in MS is by itself a challenge (49) and the absolute safetiness of OCT, together with its growing implementation in monitoring MS patients for other purposes, represents a major opportunity for current and future developments in the field.

The hope raised after the first report showing positive effects in clinical trials of a compound as remyelinating agent for MS patients (8), has been temperated with the evidences pointing to the need of further remyelinating treatments to complement currently available therapeutic arsenal to treat MS with different compounds that would result in different combinations with immunemodulators for the better matching to the clinical ideosincrasy of MS patients (50). This is event more evident after recent demonstrations of molecular changes in MS impacting on the (re)myelinating scenario in the adult CNS, including changes in oligodendroglial populations (51–55). Thus, we decided to performed the study here described using the remyelinating agent VP3.15 as pharmacological tool. In agreement with previous reports performed in different experimental paradigms and models (35, 36, 38) our results showed that the treatment with VP3.15 acted since the very first moment of application *in vivo*, ameliorating the clinical score, the score fraction and the thinning of the optic nerve measured by OCT. This is the first time where measures from OCT have been correlated with neurological clinical scores in the EAE model pointing to a useful non-invasive technique to follow the pathology with great translational potential.

The anti-inflammatory *in vivo* effect of VP3.15 seems to rely on its potential to diminish reactive (ameboid) microglia and it is in agreement with previous results from primary microglia and astrocytes cell cultures (34). Microglia have traditionally been recognized as the cells that mediate the inflammatory response in the CNS. However, it is currently known that ramified microglia, also called quiescent microglia, are involved in the structural and functional plasticity of neurons and, once polarized into anti-inflammatory phenotypes, are *sine qua non* condition for debris fagocitosis and effective remyelination (5, 44, 56–60).

Once confirmed this combined remyelinating-plus-neuroprotective-plus-antiinflammatory effect of VP3.15, its potential to treat MS should be tested in clinical trials. Up to date, there are no treatments with confirmed activity on these three incontournable aspects of MS pathogenesis (9, 50, 61, 62), confirming the inhibition of phosphodiesterases and GSK-3β as a big hope for the treatment of MS (63, 64).

Altogether, our current results, besides to supporting OCT as a useful non-invasive technique for the dynamic evaluation of retinal and optic nerve changes in EAE mice, point to the correlation between demyelinating/remyelinating events in the optic nerve and the parenchymatous CNS. While the first should be enough to include OCT in the preclinical studies on the search of effective remyelinating approaches, the second could be very useful to check evolution of MS patients and the effectiveness of the treatments that they receive, nowadays and when the desired remyelinating therapies will be a reality. It is true that our present results allowed us to measure parenchymatous CNS myelin and changes in the oligodendroglial lineage only at the end of the study, but it is also true that perform OCTs to monitorize treatments in MS patients will be faster and more affordable than monitorize via sequential MRIs. Like in most of the diseases, the finding of new useful biomarkers for MS represents a major goal for current Neurology (65).

In summary, this study has confirmed the relevant pharmacological profile of VP3.15 as a disease-modifying agent for MS with a combined remyelinating-plus-neuroprotective-plus-antiinflammatory effect and more importantly has provided evidences of OCT as a valuable methodology to follow the pathology evolution *in vivo*. The translation of these results to the clinic may open a new avenue to follow the efficacy of remyelinating drugs on patients and underscore the relevance of future clinical trials for remyelinating drugs such as VP3.15.

## Material and methods

### Induction of EAE and treatment

Six-week-old female C57/BL6 mice, bred at the CIB Margarita Salas-CSIC, were induced for EAE as has been done by our research group for a decade (66–69). Mice were divided into three experimental groups as follows: EAE-VP3.15 (compound-treated EAE animals; n=10), EAE-Vehicle (non-treated EAE animals; abbreviated as EAE-Veh; n=7) and SHAM (same procedure as EAE animals except the MOG injection; n=13). After being anesthetized with xilacine and ketamine (Xilagesic® 20 mg/ml, Calier; Ketolar® 37 mg/kg, Pfizer) chronic progressive EAE was induced by subcutaneous immunization with 250 µg of myelin oligodendrocyte glycoprotein (MOG35-55 peptide: GenScrip HK Limited. Hong Kong) emulsified in complete Freund’s adjuvant (CFA) containing 4 mg of heat inactivated Mycobacterium tuberculosis (BD Biosciences, Franklin Lakes, New Jersey, USA) and at a final volume of 200 µl. Sham-operated animals received phosphate-buffer saline (PBS) instead of the MOG peptide. Both immunized and SHAM mice were administered with Pertussis toxin (400 ng/mouse, Sigma-Aldrich, St. Louis, MO, USA), injected intravenously through the tail vein on the day of immunization and 48 h later. EAE was scored clinically on a daily basis in a double-blind manner as follows: 0, no detectable signs of EAE; 1, paralyzed tail; 2, weakness or unilateral partial hindlimb paralysis; 3 complete bilateral hindlimb paralysis; 4, total paralysis of forelimbs and hindlimbs; and 5, death. We established the peak of the symptoms as the second day of maximal clinical score reached by the mouse, being higher than 2 and the average 2.5. From this point onwards, the animals classified as EAE-VP 3.15 and EAE-VEH were intraperitoneally injected a daily dose of 10 mg/kg of VP 3.15 compound, synthesized in the CIB Margarita Salas facilities following synthetic procedures previously described (34). VP3.15 was resuspended in 100 mg/mL in DMSO diluted 1:50 in a solution of 5 % Tocrisolve (Tocris®, Minneapolis, USA) to a maximal final volume of 250 µl and the treatment was during 21 days. The EAE-VEH and SHAM animals were intraperitoneally injected with the vehicle (DMSO diluted in a solution of Tocrisolve). The exponential expression of VP3.15 healing ability was calculated using XMGrace, taking the maximal clinical score (CS max) as starting point. In order to compare both groups we normalized the score at every give time point by the maximal score (S_f_; score fraction). Thus, CS/CSmax decayed from 1 to a final plateau value, that infers the remaining score at long/infinite times (*S*∞).

All experiments were performed in compliance with the “Principles of laboratory animal care” (NIH publication No. 86-23, revised 1985), the European guidelines for animal research (European Communities Council Directives 2010/63/EU, 90/219/EEC, Regulation (EC) No. 1946/2003), and with the Spanish National and Regional Guidelines for Animal Experimentation and the Use of Genetically Modified Organisms (RD 53/2013 and 178/2004, Ley 32/2007 and9/2003, Decreto 320/2010); andapproved by the institutional ethical committees (Ethics Committees at the Consejo Superior de Investigaciones Científicas and Comunidad de Madrid).

### OCT data acquisition and analysis

For OCT analysis, a Spectralis OCT set up (Heidelberg Engineering GmbH©) was used. Both eyes of the animals were analyzed at seven time points: at the baseline (previous to immunization), onset, peak, and every five days until sacrifice (Fig. 1A). The mice were anesthetized with Ketamine 37 mg/kg (Ketolar® 50mg/ml, Pfizer) and Medetomidine 10 mg/kg (Domtor® 1 mg/ml, Ecuphar) and the eye pupil was dilated with atropine 1 % (Colircusí®, Novartis Farmacéutica, S.A.) for a better visualization. The eye under analysis was protected with a contact lens and the other with saline or artificial teardrops (Artific® 3,2 mg/ml) to avoid drying. Once immobilized, the eye bottom was focused (0.5-1.5 optic diopters) and images were captured in a high resolution mode in two scanning modalities: radial (6 radius per image) and longitudinal. The automatic real time (ART) mode was set at 45 and images were captured with a quality higher than 40 %. The follow-up tool was used so the images for the mice were acquired at exactly the same area each time. Animals were awaken with Antisedan® 5 mg/ml (Ecuphar). The area, perimeter and width of the optic disc was analysed with the Image J software (Image J). For the retinal layer thickness analysis, five points separated by 50 µm were taken at both sides of the optic nerve papilla per eye. Rretinal nerve fiber layer (RNFL), ganglion cell layer (GCL) and inner plexiform layer (IP) thickness were measured alone or combined and referred to the total retina thickness using the segmentation provided by the Spectralis OCT software. As we did not find a difference between the right and left eyes, the results are reported as a mean ± SEM of both eyes taken together. Data were taken from every single time point. The baseline OCT values were taken from animals before the immunization procedure.

### Tissue section

For histology, animals were euthanized at the final point with a lethal dose of pentobarbital (EUTANAX®, Fatro Ibérica). All animals were perfused transcardially with 4% paraformaldehyde in 0.1 M Phosphate Buffer (PB, pH 7.4). The spinal cords and optic nerves were dissected out and post-fixed in the same fixative for 4 h at room temperature (RT). After a progressive immersion in 10 %, 20 % and 30 % (w/v) sucrose diluted in 0.1 M PB (pH 7.4) for 12 h, coronal cryostat sections (20 µm thick: Leica, Nussloch, Germany) were thaw-mounted on Superfrost®Plus slides with an embedding medium for frozen samples named OCT (Tissue-Tek)®. In the case of the optic nerve, the medium used was blue TFM(tm) (Tissue Freezing Medium, Electron Microscopy Sciences, VWR).

### Eriochrome cyanine staining

For myelin analysis, the sections were dried for 2 h at RT and for 1.5 h at 37°C in a slide warmer. The slides were then placed in a container with acetone for 5 min at RT and air-dried for 30 min. The sections were stained in 0.2 % eriochrome cyanine (EC) solution for 30 min and differentiated in 5 % iron aluminium and borax-ferricyanide for 10 and 5 min, respectively, briefly rinsing under tap and distilled water between each step. Myelin is stained in blue and cell bodies in whitish (66).

### Immunohistochemistry

Sections were first air-dried for 1 h at RT. After several rinses with PB, the sections were pre-treated for 15 min with 10 % methanol in PB and washed three times with PBS for 10 min each time. In the case of myelin basic protein (MBP) labelling, sections were delipidated with a battery of ethanol-increasing concentrations (25, 75, 95 and 100 %) and later step-back rehydration for the following steps. Sections were then pre-incubated for 1 h at RT in incubation buffer: 5 % normal donkey serum (EMD Millipore, USA) and 0.2 % Triton X-100 (Sigma-Aldrich) diluted in PBS. Immunohistochemistry was performed by incubating the sections overnight at 4 °C with the primary antibodies (Table 1) diluted in incubation buffer. After rinsing, the sections were incubated with the corresponding fluorescent (1:1000, Invitrogen, Paisley, UK) secondary antibodies in the incubation buffer for 1 h at RT. In all cases, cell nuclei were stained with Hoeschst 33342 (10 µg/mL, Sigma-Aldrich) and the sections were mounted with coverslips in Fluoromount-G (Southern Biotech, Birmingham, AL, USA).

**Table 1.** List of antibodies used in this study.

| Antibody | Target | Cellular location | Dilution | Host species | Class | Manufacturer | Antibody ID |
| --- | --- | --- | --- | --- | --- | --- | --- |
| **MBP** | Myelin | Plasma Membrane | 1:500 | Rat | Monoclonal clone 12 | Biorad | aa 82--87 |
| **NFH** | Neurons/ Axons | Cell Body | 1:1000 | Rabbit | Polyclonal | Abcam | Ab 8135 |
| **Iba-1** | Microglia | Plasma membrane | 1:500 | Guinea pig | Polyclonal | Synaptic Systems | 234 004 |
| **PDGFRα** | OPCs | Plasma membrane | 1:200 | Goat | Polyclonal | RD Systems | AF 1062 |
| **CC1** | Mature oligodendrocytes | Cell body | 1:200 | Mouse | Monoclonal clone CC1 | Merck Millipore | OP 80 |
| **Olig2** | Oligodendrocyte lineage | nucleus | 1:200 | Rabbit | Polyclonal | Merck Millipore | AB 9610 |

Abbreviations: MBP= myelin binding protein; NFH=neurofilament heavy polypeptide; PDGFRα = platelet-derived growth factor receptor $\alpha$ .

### Tissue analysis and cell quantification

Eriochrome cyanine-stained sections were analyzed by acquiring 20x reconstructed images with a color camera of a Zeiss microscope (Leica DM 750) in the Microscopy Facility of Cajal Institute-CSIC and the total number of lesions was quantified using the Image J software. For all immunohistochemistry, pictures were acquired with a confocal SP5 microscope (Leica) located at the Microscopy Service of Cajal Institute-CSIC (20 µm z-stack at 3 µm intervals, 40x objective; and for microglia, 20 µm z-stack at 0.5 µm intervals, 63x objective) and cells were counted avoiding their overlap. The myelin basic protein (MBP) and the neurofilament heavy (NFH) labelling were quantified by measuring the fluorescence area using the Image J software. The total number of cells within the CNS tissue was assessed using Microscopy Image Analysis software (IMARIS, Oxford, UK), counting cells in 3 sections of spinal cord from each animal, under a user-defined threshold of fluorescence. In the case of the oligodendrocytes, we quantified the PDGFRα^+^Olig2^+^ and the CC1^+^Olig2^+^. Total Iba-1^+^ cells were sorted out into three morphological subtypes (ramified microglia, stellate-shaped microglia and amoeboid microglia cells) as detailed by Torres-Platas et al. (2014), Lee et al. (2018) and Parra et al (2019). Briefly, ramified microglia has a small and rounded cell body with processes in the form of tree branches; the stellate-shaped microglia have a large oval cell body with less extensive and thick processes; and the amoeboid microglia does not have a completely rounded cell body and its processes are not as well-defined.

### Statistical analysis

The data were expressed as the mean ± SEM and analyzed with Sigma Plot version 11.0 (Systat Software, San Jose, CA, USA). Student’s *t*-test was used to compare pairs of the different groups of mice with a Mann-Whitney U test for non-parametric data. A two way ANOVA with multiple comparisons using a Bonferroni post-hoc testwas used for the comparison of three groups (EAE-VP3.15 *vs*. EAE-VEH *vs*. SHAM), obtaining the area under the curve. Correlation analysis were performed using the Pearson’s correlation test. Minimal statistical significance was set at p< 0.05: * or # p< 0.05, ** or ## p<0.01; ***or ### p<0.001.

## Abbreviations

ART: automatic real time
CFA: complete Freund’s adjuvant
CNS: central nervous system
EAE: experimental autoimmune encephalomyelitis
EC: eriochrome cyanine
FGF: fibroblast growth factor
GCL: ganglion cell layer
IPL: inner plexiform layer
MBP: myelin basic protein
MOG: myelin oligodendrocyte glycoprotein
MRI: magnetic resonance imaging
MS: multiple sclerosis
NFH: neurofilament heavy
NMO: neuromyelitis optica
OCT: optic coherence tomography
ON: optic neuritis
PDE7: phosphodiesterase 7
PET: Positron Emission Tomography
RNFL: retinal nerve fiber layer
RXR: retinoid X receptor
VDR: vitamin D receptor
VEH: vehicle

## Acknowledgements

Funding from Asociación Española de Esclerosis Múltipe-Confederación Española de Personas con Discapacidad Física y Orgánica (AEDEM-COCEMFE; Dña. Angela Sastre legacy) and Ministerio de Ciencia, innovación y Universidades (grant no. SAF2015-72325-EXP, SAF2016-77575-R to FdC and RD16-0019 to FdC) is acknowledged.

The authors would like to thank the invaluable help of Sergio Casas from Instituto Cajal with the IMARIS equipment, the animal facility at CIB Margarita Salas for their technical assistance and support, Eduardo Sanz from Universidad Complutense for the exponential adjustment and Carmen Hernández and Belén García from the Confocal Microscopy facility at Instituto Cajal for their help in image acquisition.

